# The yEvo Mutation Browser: Enhancing student understanding of experimental evolution and genomics through interactive data visualization

**DOI:** 10.1101/2025.07.18.665463

**Authors:** Leah Anderson, Jasmine Schoch, Eleftheria Anastasia, Faheem Azeemi, Zilong Zeng, Virginia Wang, Sayeh Gorjifard, Adrianna Matos-Nieves, Jordan Safer, Sumaiya Iqbal, Maitreya J. Dunham

## Abstract

Experimental evolution is a powerful method for studying the relationship between genotype and phenotype by observing how populations genetically adapt to controlled selective pressures. In educational settings, this approach also offers a dynamic way for students to engage with molecular genetics. One such educational effort, known as “yEvo” (<u>y</u>east <u>evo</u>lution), introduces experimental evolution into high school classrooms, allowing students to evolve the baker’s yeast *Saccharomyces cerevisiae* under various stressors and investigate the resulting genetic changes. While the hands-on experiments have been successful in fostering student interest and understanding of evolution, the downstream data analysis (interpreting whole-genome sequencing results of evolved yeast compared to the ancestor) remains a challenge. Students often struggle to grasp the significance of their mutated genes and lack the broader context to determine which mutations are most phenotypically relevant. To address these issues, we developed the yEvo Mutation Browser, an intuitive web tool designed to assist students and researchers alike in visualizing and contextualizing genome sequencing data. Developed using R Shiny, this tool features an interactive chromosome map displaying mutated genes, graphs categorizing mutation types, a gene viewer illustrating specific mutation sites within genes, and a protein view that maps specific mutations onto protein structures. The app also features an option for non-yEvo-affiliated users to upload their own experimental evolution or genetic screen datasets and compare them with all yEvo data collected since 2018. The yEvo Mutation Browser streamlines data interpretation, helping students understand how organisms employ diverse genetic strategies to adapt to environmental stress. In the future, this framework could be adapted for use with other model organisms, offering a valuable resource for both genetics research and education.

**Significance:** Bringing novel research into the classroom can transform how students learn science. The yEvo project engages high school students in experimental evolution, enabling them to witness natural selection in action and connect genetic mutations to evolutionary outcomes. By providing an accessible web-based tool, the “yEvo Mutation Browser,” this work lowers barriers to interpreting genome sequencing data, allowing students, teachers, and researchers alike to explore how yeast populations adapt to diverse stress conditions. This resource not only strengthens STEM education by making cutting-edge genetics more approachable, but also builds a shared dataset that enriches the broader yeast genetics community.

## Introduction

Experimental evolution is a powerful tool for studying the genotype-phenotype relationship by subjecting populations to selective pressures that drive adaptation over time via genetic mutations. In microbiology, experimental evolution is conducted by propagating microbes under suboptimal conditions such as nutrient limitation, drug exposure, or chemical stress, which restricts their growth. Under these selective conditions, rare spontaneous mutations that improve survival and growth become enriched in the population. These mutations can then be detected through whole genome sequencing, allowing researchers to pinpoint genetic changes associated with the selected trait. Follow-up experiments can then verify whether these mutations are causative for the observed phenotypic changes. This approach is not only a robust method for studying molecular adaptation but also provides a broader understanding of the role of genetic variation on meaningful phenotypic changes. In contrast to classical genetic screens, which are typically designed to find single, large effect mutations, experimental evolution elucidates the combined effect of multiple mutations with quantitative effects.

Experimental evolution is a valuable tool not only for research but also for education, offering a hands-on approach to understanding key concepts in evolution and genetics (Bennett *et al*. 2021; Cooper *et al*. 2019; Dahan *et al*. 2019; Ratcliff *et al*. 2014; Smith *et al*. 2016; Taylor *et al*. 2024; Matela *et al*. 2025). One example of this is the “yEvo” (<u>y</u>east <u>evo</u>lution) project, a series of laboratory modules designed for high school biology students to experimentally evolve the baker’s yeast *Saccharomyces cerevisiae*. Through this approach, students gain practical laboratory skills, such as sterile technique and liquid handling, while directly observing evolutionary processes in action. The yEvo curriculum deepens students’ understanding of microbial evolution, genetic variation, and the genetic basis of adaptation, while also boosting confidence in designing experiments and fostering interest in STEM careers (Taylor and Warwick *et al*. 2024). In addition to its educational impact, the data generated in yEvo classrooms has contributed to published scientific studies, offering insights into yeast adaptation and demonstrating the broader applicability of this teaching model (Geck *et al*. 2024; Moresi *et al*. 2023; Taylor *et al*. 2022). The success of yEvo lies in its use of *S. cerevisiae* as an ideal classroom model organism. Yeast is easy to grow, has a well-characterized genome, and adapts to new environments in just a few weeks. Students work in lab groups to expose yeast populations to stressors such as clotrimazole, caffeine, or other agents, simulating real-world evolutionary pressures like exposure to antimicrobial drugs. After the experiments, samples are analyzed for genomic mutations, and students use resources like the *Saccharomyces* Genome Database (SGD) (Engel *et al*. 2024) to hypothesize how specific mutations may drive adaptation.

Despite the success of the hands-on experimental evolution portion of the yEvo project, the downstream analysis and hypothesis-building steps have proven to be more challenging for students. Many students report that the final stages, which involve interpreting whole genome sequencing data and making connections between their mutated gene lists and the stress conditions, feel much less interactive and engaging than the evolution experiments themselves. Historically, our mutation data analysis module had students review lists of mutated genes from their lab groups’ samples and conduct independent online research using resources like SGD and primary literature. However, through this approach, students often found it difficult to understand the significance of their mutated genes. Additionally, students were lacking the broader context to determine which of their identified genes are most likely to be causative for the observed phenotypic changes, as they only have access to their group’s data without knowing how reproducible their results might be. To keep students engaged through the entirety of the project and help them see the big picture, there has been a need for a user-friendly, visually appealing approach to data analysis that makes complex genomic data easier to interpret.

To meet this need, we have developed the “yEvo Mutation Browser” (available at yevo.org/mutation-browser). This web-browser based application (web app) was written using the R Shiny package, and offers an accessible and engaging way for students to interact with their data. The browser was specifically designed to be user-friendly for students while capturing the full scope of their experimental evolution projects, including linking them to the larger yEvo network of classrooms. However, the tool also allows users to upload their own data locally, expanding its possible use to researchers such as yeast geneticists, who would like to interact with their own experimental evolution or genetic screen data. We have made the code available such that other users could also create their own instances of the browser populated with the data of their choice. Here, we outline the key features and applications of the yEvo Mutation Browser.

## Methods

### Classroom experiments and sequencing

Classroom evolution protocols, sample preparation, and sequencing were performed as previously described (Geck et al 2024, Moresi et al 2023, Taylor et al 2022, Taylor and Warwick *et al*. 2024). In high school classrooms, students evolved *Saccharomyces cerevisiae* under various selective pressures including caffeine, clotrimazole, echinocandins, acetic acid, sodium bicarbonate, wood hydrolysate, and tea tree oil. After the experimental evolution phase, university collaborators isolated three individual yeast clones from each lab group’s evolved population. One clone per group showing a clear phenotypic improvement was selected for Illumina whole-genome sequencing. The sequencing data were analyzed using a custom variant-calling computational pipeline (available at: https://github.com/leahmarieanderson/exp_evo_variant_calling). Sequencing reads were aligned to the *S. cerevisiae* S288C reference genome using Burrows-Wheeler Aligner (BWA) (Li and Durbin 2009). Mutations were called using three variant callers: gatk4 (McKenna *et al*. 2010), lofreq (Wilm *et al*. 2012), and freebayes (Garrison and Marth 2012). Quality filters were applied to the variant calling files (VCFs) via bcftools (Danacek *et al*. 2021), and ancestor variants were removed from evolved VCFs using bedtools (Quinlan and Hall 2010). Variant annotations were then added via a custom annotation python script (also available at the git repository above). Transposon insertions were called using the McClintock package (Nelson, *et al*. 2017). Annotated variants include coding mutations (such as nonsense, missense, synonymous, frameshift), non-coding mutations (such as intergenic, intronic, non-coding RNA), and transposon insertions. Variants within 200 base pairs upstream of a start codon were labeled 5’ upstream mutations. All mutations passing the quality thresholds were manually reviewed in the Integrative Genomics Viewer (IGV) (Robinson *et al*. 2017) to confirm that the variant was both a high quality call and present only in the evolved sample.

### yEvo Mutation Browser R Shiny app

The yEvo Mutation Browser was developed in R and leverages the Shiny package to build the app’s user interface (UI). The plotly package was used to create interactive visualizations either as a wrapper around a static ggplot graph or implemented directly. The Shiny web app is deployed on the shinyapps.io platform. When a user accesses the yEvo Mutation Browser, a new session is spun up, loading a comma-separated values (CSV) file storing the yEvo Cumulative Database, which contains all yEvo mutation data dating back to 2018. These data were bundled into the web app during deployment to shinyapps.io, and each new session loads its own isolated copy of the data. The Shiny framework assigns each user a unique session, enabling multiple users to explore the app concurrently without interference. Within a session, users can interactively filter the default dataset or upload their own mutation data in CSV format; uploaded data is then appended only to that user’s session-specific copy and exists solely in memory for the duration of that session. These temporary uploads do not modify the original yEvo Cumulative Database or affect other users’ sessions.

The mutation browser’s UI is split into 2 main sections: The filtering panel and the visualization panel. The filtering panel allows the user to upload their own data and select key attributes to narrow the points being visualized for more thorough analysis and customization. At the top of the visualization panel, the user can select their desired method (Chromosome Map, Variant Pie Chart, SNP Counts, Gene View, or Protein View) to view the filtered data. The filtered data can also be viewed under the “Table” tab, and downloaded in CSV format.

### Protein View tab

Yeast gene names and associated data, including SGD identifiers, chromosome numbers, biological pathways, and genomic coordinates, were retrieved using the AllianceMine API (Mistry *et al*., 2020). These data provide a genomic backbone for cross-referencing with protein information. Each yeast gene is mapped to its corresponding Uniprot ID via the UniProt Mapping API (The Uniprot Consortium, 2025), resulting in 6,823 mappable proteins out of 6,909 annotated yeast genes. Providing MolStar with these Uniprot identifiers allows it to render the canonical AlphaFold-predicted 3D protein structures. To enrich structural interpretation, motif and domain annotations are collected from the UniProt REST API using UniParc-linked entries, enabling access to PFAM (The Alliance of Genome Resources Consortium, 2024) domain annotations while maintaining consistent sequencing tracking across databases.

Protein structures were rendered via the *Mol (MolStar) API\**, a web-based molecular visualization engine optimized for 3D structural analysis (Sehnal *et al*., 2021). Integration into R Shiny required modular bundling from Webpack, allowing incorporation of custom JavaScript functionality. This allows for further interaction between MolStar and its users, by providing the ability to dynamically highlight residues, domains, and motifs; overlay mutations onto structure; control zoom and rotation; and give structural context visualization for uploaded variants.

### Software availability and requirements

The yEvo Mutation Browser can be accessed via Shiny apps (https://leahmarieanderson.shinyapps.io/yEvoMutBrowser/) or on the yEvo website (https://yevo.org/mutation-browser/). The web app can also be installed and run locally as per the instructions provided on the GitHub repository.

The code base for our app is freely available on GitHub at https://github.com/dunhamlab/yEvoMutBrowser as an R application. The installation requires R (>= 4.3.0). We have tested local versions on both macOS and Windows operating systems. Please refer to the Github repository for more information on installation.

### Large language model usage

Claude Code (Sonnet 4, Anthropic) and ChatGPT (versions 4o and 4.5, OpenAI) were used as supportive tools during the development of the yEvo Mutation Browser. These tools assisted with refining our existing code by suggesting methods and formatting to improve efficiency and readability. ChatGPT was also consulted during manuscript preparation for language editing and clarity suggestions. All AI suggestions were carefully reviewed by the authors after implementation to ensure accuracy and integrity of the manuscript and codebase.

## Results

### yEvo Mutation Browser usage and visualizations

The yEvo Mutation Browser is built using R Shiny (https://shiny.posit.co/), featuring a user-friendly interface that allows users to toggle between different data visualization options (genome location, variant type, gene, protein, etc.), while the underlying server processes and displays the selected information reactively. This functionality enables users to explore various datasets, generating visualizations tailored to their specific selections. No tool previously existed for our students to visualize their own data, and the Mutation Browser now fills a major gap in the broader yEvo project. The Mutation Browser was used in all 2024-2025 yEvo classrooms, with teacher and student feedback incorporated into its design. To date, there are over 2000 genomic variants included in this dataset, and more will be added as classrooms continue to participate in yEvo. The source code and datasets used to develop the yEvo Mutation Browser are publicly available at https://github.com/dunhamlab/yEvoMutBrowser. In the following sections, we describe the application’s usage, its user interface, and the visualizations users encounter while navigating the app.

#### Application usage

Upon launch, all cumulative yEvo data collected since 2018 is loaded into memory. This data exists in a comma-separated-values (CSV) file which is then used to generate all interactive visualizations (see Methods). Users begin by selecting the data subset they wish to explore. One option is to view data by classroom, where users can choose either the entire class dataset or individual lab group data using a dropdown menu after specifying their class (Figure S1A); this is particularly useful for students who have completed the experimental evolution module of yEvo with their specific classroom. Alternatively, users can select “View by Selection Condition,” which displays all the cumulative yEvo data generated under specific selection conditions (Figure S1B). Users are able to see all the selection conditions used in yEvo since 2018: the antifungal agents clotrimazole, caspofungin, micafungin, and tea tree oil, and the industrial/environmental conditions acetic acid, caffeine, sodium bicarbonate, and wood hydrolysate. While most data in the app is from experimental evolution of the haploid lab strain S288C, some experiments began with different backgrounds, including a diploid version of S288C, a baking strain, or genetically modified knockout strains. If the selected dataset contains multiple strain options, users can specify which strain(s) they want to view using the “Ancestor Strain” dropdown menu (Figure S1C and D). For simplicity, results from all strains are viewed against the S288C reference genome, though this could be modified in the future (see discussion).

The app also allows users to upload their own Variant Call Format (VCF) files in CSV format, which enables any user to visualize their own data (see Methods). Instructions for uploading datasets are provided in both the “Tutorial” tab within the app and on the README on our GitHub repository. When uploading, users are prompted to input their name and year, which then become accessible via the “View by Class” dropdown menu (Figure S1E). This new data is then temporarily appended to the yEvo dataset available on the app for that user’s session. If the selection condition in the uploaded dataset matches pre-existing yEvo data on the server, the new data is added to the cumulative dataset for that condition, allowing users to see how their results compare with patterns observed in our yEvo experiments. This feature is beneficial not only for educators but also for researchers seeking a comparison of their data with existing experimental evolution datasets.

Once the desired dataset is selected, users can access all of the available data visualizations. Users can also modify their dataset selections while viewing any of the visual plots, which are described in the following sections.

#### Visualizations

##### 1. Chromosome plot

The first visualization users encounter in the yEvo Mutation Browser is the chromosome plot, which provides a linear representation of the 16 nuclear chromosomes along with the mitochondrial genome of *S. cerevisiae* (Figure 1A). This plot displays the locations of all mutated genes in the selected dataset, marked at their chromosomal positions. For simplicity of viewing, only mutations in coding regions are shown on the chromosome plot. Tick marks along the linear chromosomes indicate these sites, and hovering the mouse over each tick mark reveals the name of the mutated gene, as well as how many times it was mutated in repeat experiments, which is also indicated visually using a color scale. Users can also zoom in on specific regions by clicking and dragging, offering a closer look at mutations in a given area (Figure S2). Clicking on a gene in the chromosome plot will take the user to “Gene view” for that gene, where they can explore precise locations of mutations in that gene (see Gene view description below).

**Figure 1:**
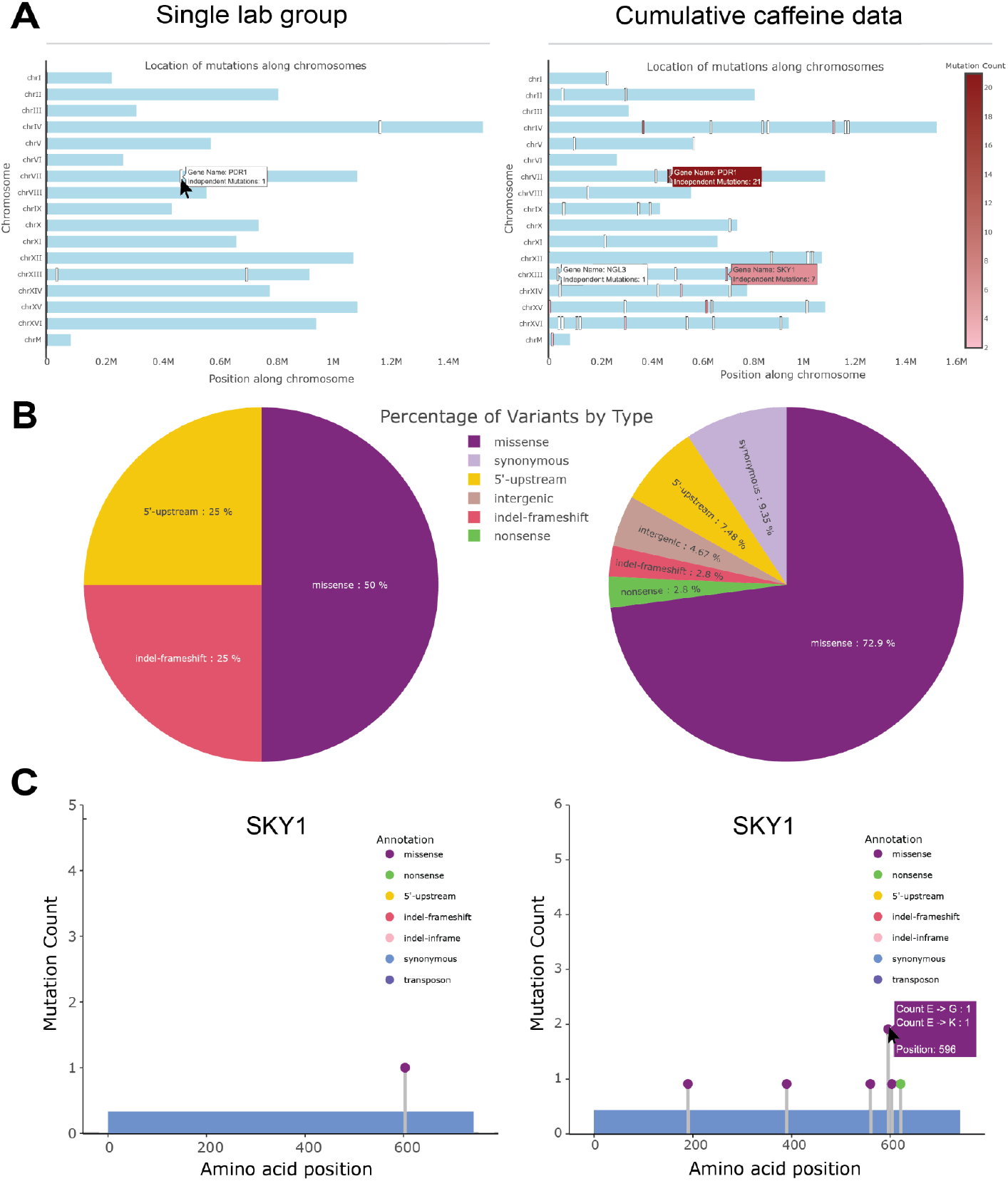
Visualization examples of a single lab group vs cumulative classroom data. Visualizations of data from the Brewer 2022 classroom, where yeast was evolved in caffeine. (A) Chromosome plot showing genes mutated in a single sample (left) or the whole class (right). The scale bar on the right shows the number of samples that had a mutation in that gene. (B) Pie chart showing makeup of mutation types. (C) Gene view displaying where each coding mutation affected the gene by amino acid position. Color indicates the type of mutation.

The chromosome plot allows students to directly compare their results with those from other students within their own classrooms as well as to previous years and other yEvo sites performing similar experiments, providing a more robust and visually intuitive way to assess the significance of genetic mutations. Figure 1A highlights this feature. The left panel shows data from a single lab group who evolved yeast in increasing concentrations of caffeine, where four mutated genes are mapped: *SVF1, PDR1, NGL3*, and *SKY1*. Once a user switches the view to “View by Condition” and selects “caffeine,” the chromosome map is labeled with how recurrently each gene in the dataset was mutated in the cumulative caffeine yEvo data, with a white-to-red gradient displaying frequency (Figure 1A, right panel). Using the “Selection condition” view, it can be seen that *PDR1* and *SKY1* were mutated in 21 and 7 samples respectively, while *NGL3* and *SVF1* were unique to the single sample. This reinforces the likelihood that *PDR1* and *SKY1* play a key role in adaptation to caffeine, which has now been experimentally verified since the high school experiments were carried out (Geck *et al*. 2024).

##### 2. Pie chart of mutation types

The second visualization provided by the yEvo Mutation Browser is a pie chart that displays the distribution of mutation types identified by the sequencing analysis pipeline. Unlike the chromosome plot, which focuses solely on the locations of mutated genes, the pie chart encompasses all variants, including those in non-coding regions (such as intergenic, intronic, tRNA, rRNA, noncoding RNA, and autonomously replicating sequence regions). This gives students a broader overview of the types of genetic changes that occurred in their evolution experiments. One key insight students gain from this visualization is that, regardless of the selection condition, the majority of mutations in the evolved samples are those that alter protein sequences. For instance, the left panel of Figure 1B shows the mutation types for one lab group’s data, while the right panel shows cumulative data from multiple classrooms that evolved yeast in caffeine. In both cases, the vast majority of mutations are missense mutations. This becomes a valuable teaching tool, helping students understand that while mutations occur randomly across the genome, those that lead to phenotypic changes are primarily found in coding sequences, which are then selected for during the course of the experiment.

##### 3. Mutation spectrum

The third visualization offered by the yEvo Mutation Browser is the mutation spectrum, depicted in Figure 3B. This plot illustrates the distribution of single nucleotide polymorphisms (SNPs) and small insertions and deletions (indels). By categorizing the different types of SNPs, the mutation spectrum provides insights into the underlying biochemical processes that drive DNA mutations. Students can observe that transitions—mutations where a purine is replaced with another purine (A to G / G to A) or a pyrimidine with another pyrimidine (C to T / T to C)—are more frequent than transversions, which involve purine-to-pyrimidine exchanges and vice versa. This plot displays that SNPs and indels can occur at different frequencies during evolution, and that different mutagens or background genotypes can cause altered mutation spectra compared to the natural mutation rate. Recognizing the prevalence of transitions over transversions reinforces the concept that certain mutations are inherently more likely to occur based on the chemical stability and replication mechanisms of DNA. Further applications of this visualization, such as using it to explore how yeast strains with different mutation rates exhibit distinct mutation spectra, will be described later in this paper.

##### 4. Gene view

The fourth visualization is “Gene view,” a linear lollipop plot that displays the position of mutations along the protein product and 200 bp upstream of the start codon for each mutated gene (Figure 1C). In this view, users can select a mutated gene from a dropdown menu based on the selected dataset. Once a gene is selected, the plot reveals the length of the amino acid sequence and color-coded lollipops indicating where mutations occurred. The colors differentiate between mutation types, such as missense, nonsense, frameshift, synonymous, transposon insertions, or 5’-upstream mutations. The height of each lollipop reflects how frequently that particular site in the protein was mutated within the dataset. For example, the left panel of Figure 1C shows the gene view for an evolved yeast strain, and in the live use of the app, a dropdown menu below the gene plot would display the four genes that were mutated in that sample. In contrast, the right panel illustrates the cumulative caffeine data, with all genes mutated in that condition in the dropdown menu and the lollipop heights showing the recurrence of mutations at specific sites. Additionally, hovering the mouse over a lollipop provides information about the mutation, including the amino acid site number, residue type, and the resulting amino acid caused by the DNA mutation (if any).

Furthermore, the gene view includes a “Learn about gene” button, which redirects users to the SGD locus page for the selected gene. This feature allows students to explore the gene’s function in yeast and use this information to generate hypotheses about how mutations in that gene might contribute to observed phenotypic changes. Overall, the gene view visualization is designed to help students better conceptualize where mutations occur within genes and to encourage critical thinking about how and why certain regions may be more frequently mutated. By connecting mutation sites to their potential impact on protein function, students gain deeper insights into the molecular mechanisms driving adaptation in their experiments.

##### 5. Protein view

The final visualization is “Protein view”, inspired by the Broad Institute’s Genomics to Proteins (G2P) portal (Kwon *et al*., 2024). Similar to Gene view, users are shown a dropdown menu of the mutated genes in the selected data. Upon selection, the app renders the predicted 3D structure of that gene’s protein product. Below the protein visualization, a linear track displays amino acid position along the x axis, plotting protein domains, motifs, as well as mutation sites that are color-coded by type in accordance with the Gene view plot (Figure S3). Users can click any mutation to highlight its location, or choose “View all mutations” to project all mutations from the selected dataset (e.g. an entire classroom cohort or cumulative selection condition) onto the protein (Figure 2, right panel). Protein view is a proteome-wide interactive resource, allowing users to explore how specific mutations may affect protein stability, folding, or functional domains. This interactive visualization facilitates identification of potential clustering of mutations in 3D space and supports interpretation of likely effects on protein function.

**Figure 2:**
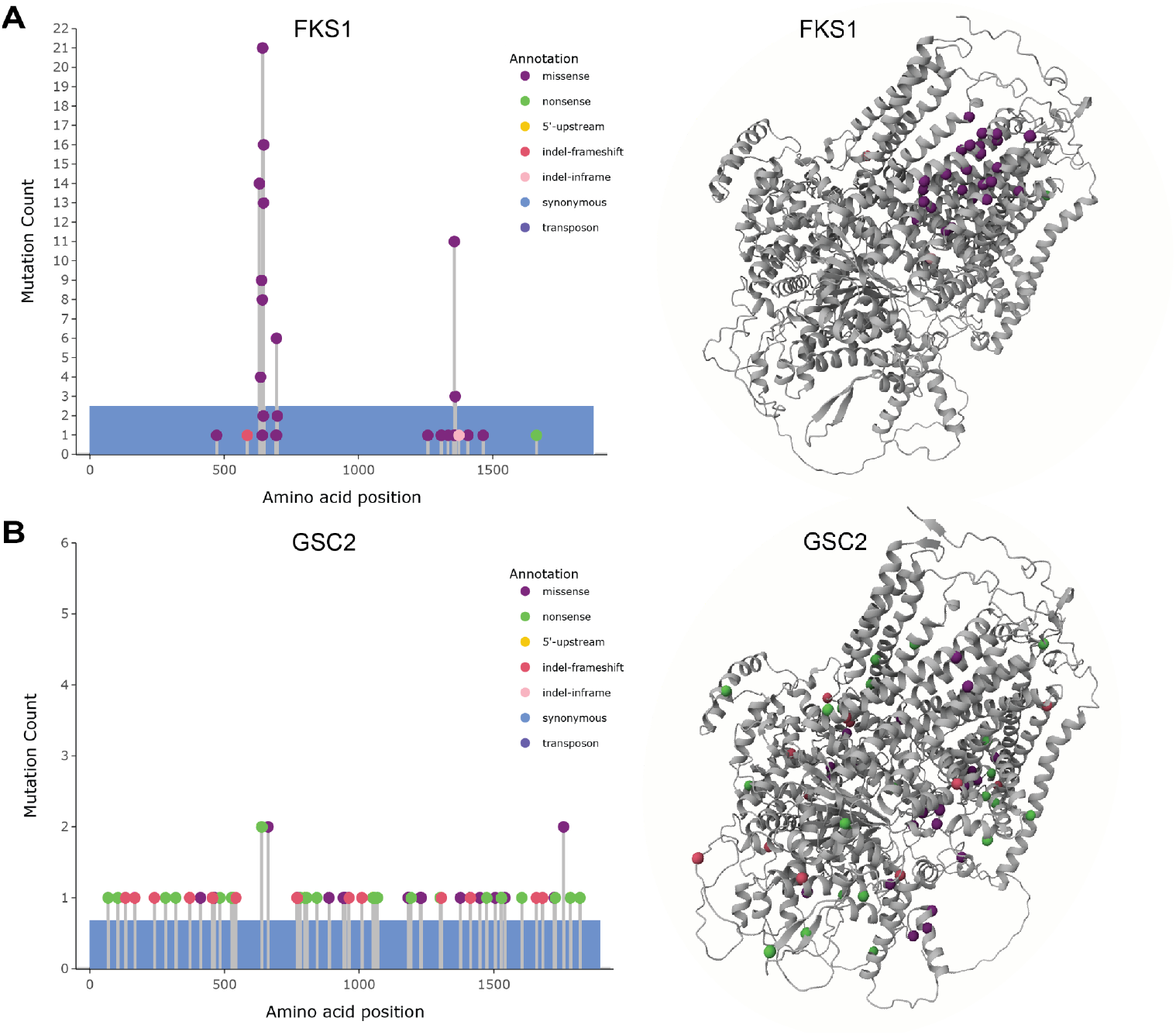
Gene view can reveal mutation patterns for specific genes. Mutations in *FKS1* (A) and *GSC2* (B) display different patterns for yeast samples evolved in caspofungin. Left panels display the location and frequency of mutations along the linear amino acid sequence of each gene, and right panels display locations of these mutations in the predicted 3D protein structure. Mutations are color coded by type, and follow the same scheme in the linear and 3D diagrams.

### Further educational applications

While we have outlined the foundational educational use cases for the yEvo Mutation Browser, there are additional benefits that can offer valuable teaching points for more advanced biology classrooms.

#### Signatures of mutation effects

As yEvo students explore gene functions and hypothesize which mutations might be phenotypically causative, predicting how specific mutations alter gene function remains a challenge. Does increased survival arise from the loss of gene function, or from a gain or alteration of function? Historically, students could only make educated guesses based on the mutation types in their genes of interest (synonymous, missense, nonsense, etc). With the yEvo Mutation Browser, students can now make more informed predictions about the nature of mutations by analyzing mutation patterns from cumulative data. For instance, when viewing cumulative data in Gene View for a particular condition, a student might observe that mutations in a particular gene are clustered in one region and are exclusively missense. A tight clustering of mutations at a single site, especially if only missense, suggests a potential gain-of-function (GOF) mutation, as there are limited ways to generate a GOF mutation that enhances fitness. Conversely, if a gene exhibits mutations scattered throughout, with a mix of missense and nonsense mutations, this pattern likely indicates a loss-of-function (LOF) mutation, as many different types of mutations can disrupt gene function. These mutation patterns have been extensively studied and are seen in various human diseases, including cancer (Gerasimavicius *et al*. 2022; Prior *et al*. 2012; Sivley *et al*. 2018; Stehr *et al*. 2011; Turner *et al*. 2015). This offers teachers a valuable opportunity to connect the mutation patterns observed in the classroom to real-world medical applications, such as cancer and other genetic disorders, making the lessons more relatable and impactful for students.

An example of these mutation signatures is depicted in Figure 2, which illustrates cumulative data for caspofungin. Students may notice using Chromosome View that two genes are particularly enriched for mutations, *FKS1* and *GSC2*. In Gene View, *FKS1* shows mutations clustered at two distinct locations, all of which are missense (Figure 2A). This suggests that mutations in *FKS1* may lead to a gain or alteration of gene function, a pattern consistent with prior research in both pathogenic yeast and *Saccharomyces* species (Katiyar and Edlind 2009; Johnson and Edlind 2012; Katiyar *et al*. 2006; Laverdiere *et al*. 2006; Park *et al*. 2005; Thompson *et al*. 2008; Balashov *et al*. 2006; Cleary *et al*. 2008; Garcia-Effron *et al*. 2008; Khan *et al*. 2007; Johnson *et al*. 2011; Perlin 2007). These missense mutations also appear to cluster together in the protein structure in regions that exist on the outside of the cell membrane, suggesting that changes in this specific location can lead to increased echinocandin resistance. In contrast, *GSC2* shows mutations spread across the gene, with many being nonsense and frameshift mutations (Figure 2B), suggesting that loss of function may contribute to the observed phenotype. In fact, this aligns with prior work which has shown that *gsc2Δ* can decrease echinocandin sensitivity in *S. cerevisiae* (Lesage *et al*. 2004). Armed with this information, students can generate more informed hypotheses. They may discover through SGD and primary literature that *FKS1* and *GSC2* are homologous genes involved in β-1,3-glucan synthesis, a critical component of fungal cell wall integrity. Moreover, they may learn that caspofungin targets the fungal cell wall by inhibiting the enzyme β-1,3-glucan synthase (Denning 2002), which is composed of catalytic subunits encoded by *FKS1* and *GSC2* (Lesage and Bussey 2006). From this, students could hypothesize that a GOF mutation in *FKS1* may alter the enzyme’s structure in a way that reduces the binding affinity of caspofungin, allowing the enzyme to retain functionality. In the same vein, loss of *GSC2* function (which the app shows almost always co-occurs with an *FKS1* missense mutation) could enable the yeast to prioritize *FKS1*-based glucan synthesis, providing resistance to caspofungin without completely impairing cell wall biosynthesis. While this idea is still hypothetical, (and we are currently exploring this hypothesis in our lab), we believe that involving students in hypothesis building on a concept that has not yet been explored can be critical for their learning and interest in science. This example also displays how leveraging the power of cumulative data visualizations in the yEvo Mutation Browser can help students think more deeply about gene function and gain a better understanding of the molecular mechanisms driving adaptive evolution.

#### Visualizing the effects of a mutator

While most yEvo classrooms conduct experimental evolution by relying on the natural mutation rate of *S. cerevisiae*, one class was interested in exploring how a higher mutation rate might affect the evolutionary process. Since many mutagenesis methods are not safe for classroom use, we provided them with a strain of *S. cerevisiae* containing a deletion of the *MSH2* gene, which encodes the Msh2 protein involved in DNA mismatch repair. Loss of *MSH2* leads to an 85-fold increase in the spontaneous mutation rate (Alani *et al*. 1995; Reenan and Kolodner 1992), and mutations in the human homolog elevate the risk of cancers (Fishel *et al*. 1993, Bridge *et al*. 2014, Win *et al*. 2017, Bowen *et al*. 2025). The class performed experimental evolution in micafungin side-by-side with both the *msh2Δ* mutator strain and the S288C lab strain that is normally used for yEvo experiments. While increasing the mutation rate might seem advantageous for rapidly observing adaptation, it can significantly complicate the sequencing data analysis. For example, instead of observing clear, repeated mutations concentrated in a handful of genes consistently linked to antifungal resistance, the mutator strain accumulates numerous additional “hitchhiker” mutations across the genome, which could be completely unrelated to the selected phenotype. Consequently, experimentally evolved mutator strains often have dozens of mutated genes, most of which do not contribute to the adaptive phenotype, thereby complicating the identification and verification of causative mutations. This concept can be challenging for students who are just learning about evolutionary biology, but the visualizations provided by the yEvo Mutation Browser clearly illustrate how employing a higher mutation rate during experimental evolution can affect the results.

To see the effects of strain backgrounds, we added a feature in the user interface that allows users to toggle between strain backgrounds when a dataset contains more than one. Therefore, when selecting the “micafungin” condition, users can choose between the S288C strain and the *msh2del* strain (Figure S1C). By comparing the chromosome plots of these two strains, users can observe that the mutator strain shows far more mutated genes, most of which are color coded white, indicating that they were only mutated in a single sample. Conversely, the S288C strain has fewer mutated genes overall, but a larger proportion of these genes were recurrently mutated across multiple samples (Figure 3A). Interestingly, in both mutator and non-mutator datasets, *FKS1* and *GSC2* remained the most frequently mutated genes. This indicates that, in this particular example, the mutator strain did not provide additional insights into new antifungal drug resistance mechanisms, as it recapitulated the same key mutations seen in the S288C control strain.

**Figure 3:**
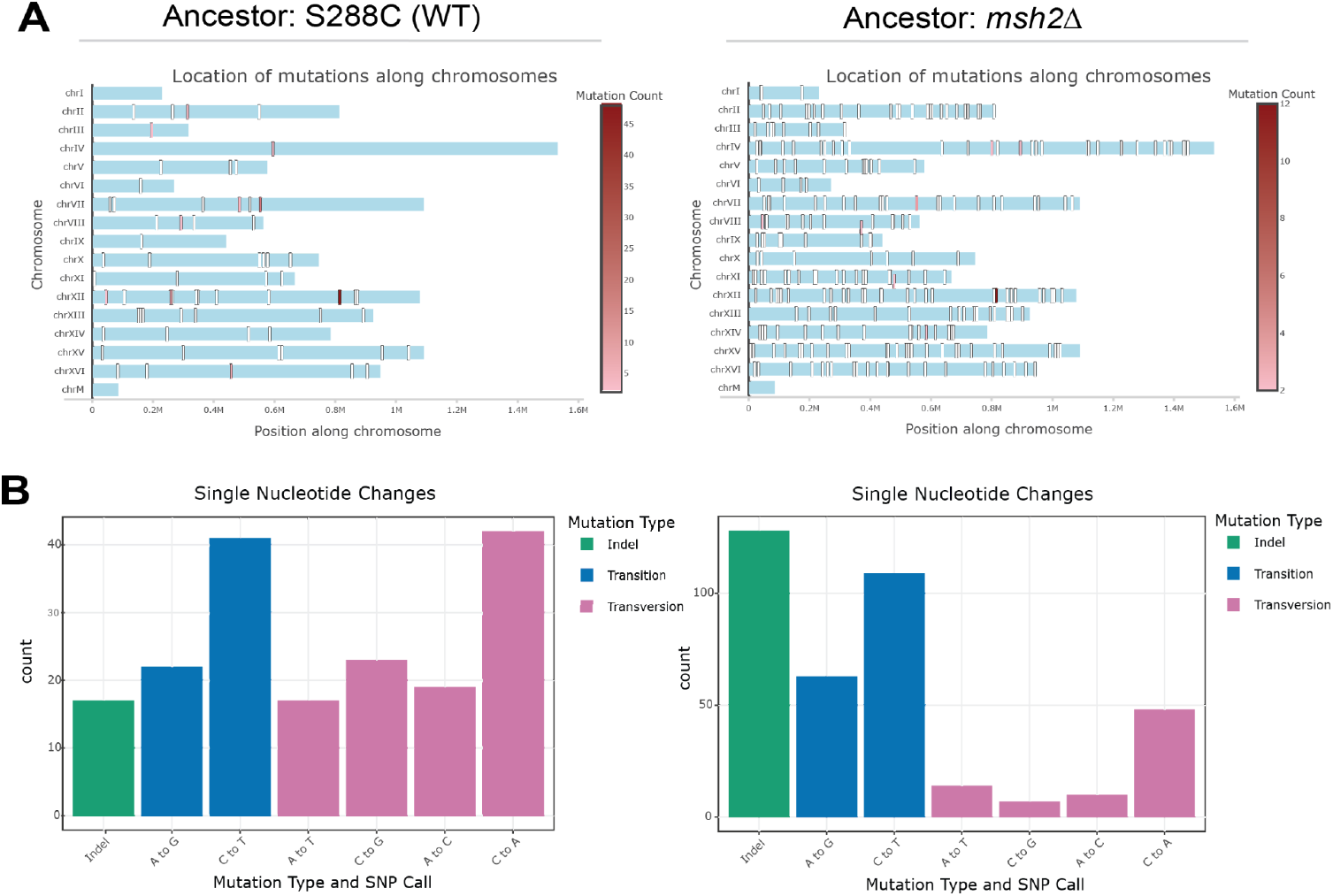
Visualizations reveal differences in mutation frequency and spectra between strains. When compared to data from an evolved S288C strain (left panels), mutations in a mutator strain missing the *MSH2* mismatch repair gene (right panels) occur in more locations across the genome (A), and show a bias towards indel mutations (B). In the ancestor strain experiments, n=37. In the deletion strain experiments, n=8.

The mutation spectrum visualization also reveals striking differences between these parallel experiments. Specifically, the mutator strain exhibits a significant bias towards indels compared to SNPs, a finding consistent with previous research (Reenan and Kolodner 1992; Alani *et al*. 1994; Greene and Jinks-Robertson, 2001; Lang *et al*. 2013). For more advanced classrooms, this provides an opportunity to delve deeper into the molecular mechanisms driving these biases, promoting critical thinking about mutation processes. Additionally, educators can use these data to reinforce the concept of hitchhiker mutations, as the mutator strain displays many more mutations across the genome, most of which are likely not significant but are carried along with beneficial mutations elsewhere in the genome. Since these results are available to all via the Mutation Browser, teachers could refer to them instead of performing the experiment themselves in order to demonstrate why relying on the natural mutation rate is sufficient for experimental evolution in yeast, while also reinforcing key concepts such as hitchhiker mutations and the complexities of mutation analysis.

## Discussion

Here we describe the yEvo Mutation Browser, an application built in R Shiny, designed to provide comparative visualizations of *S. cerevisiae* experimental evolution data. We were motivated to create this application to enhance yEvo students’ understanding of the evolutionary experiments they conducted in the classroom. Previously, our students were limited to reviewing lists of mutated genes from their lab groups’ samples and conducting independent research using data available online. With the yEvo Mutation Browser, students can now not only visualize the genomic locations of their mutations but also explore how frequently these mutations recur in repeat experiments under the same or similar conditions. This added layer of interactivity allows students to engage more deeply with their data and recognize patterns that may not have been apparent before.

Each visualization in the yEvo Mutation Browser contributes uniquely to student learning by offering insights that were not previously accessible through static gene lists. The chromosome plot, for instance, helps students better grasp the likelihood that specific genes are causative of observed phenotypes. By enabling a direct comparison of their results with those from other classrooms and experiments, students can see how certain genes are commonly mutated in repeated experiments, increasing the likelihood that mutations in these genes can affect the observed phenotype. This helps students gain broader context, which is more effective than our previous approach of having students hypothesize gene significance through isolated online searches. The pie chart visualization breaks down mutation types, helping students understand why certain variants, such as missense mutations, occur at a higher frequency. This becomes a valuable teaching tool, helping students understand that while mutations occur randomly across the genome, those that lead to phenotypic changes are primarily found in coding sequences, which are then selected for during the course of the experiment. For more advanced classrooms, the pie chart can also be used to explore why certain mutation types, such as 5’-UTR and nonsense mutations, are also frequently observed in evolved yeast samples. This also provides an opportunity to introduce students to the concept of hitchhiker mutations—those that may not directly contribute to adaptation but are carried along with beneficial mutations. This concept can be reinforced by comparing individual samples containing intergenic or synonymous mutations to their corresponding chromosome plot, helping students identify whether these mutations co-occur with coding mutations, which are more likely to be phenotypically significant. The mutation spectrum provides an opportunity to explore the biochemical mechanisms behind mutation events, distinguishing between transitions and transversions. It also can display how different mutagens or genetic backgrounds can lead to specific mutation signatures. The Gene view allows students to connect the presence of mutations with their possible functional effects, displaying “hotspot” mutation sites and the corresponding amino acid changes. This offers teachers a valuable opportunity to connect the mutation patterns observed in the classroom to real-world medical applications, such as cancer, antibiotic resistance, viral evolution, and other genetic processes, making the lessons more relatable and impactful for students.The integration of the “Learn about gene” button further streamlines the learning process by directing students to detailed information on SGD, facilitating better learning, engagement, and hypothesis development. This feature allows students to explore the gene’s function in yeast and use this information to generate hypotheses about how mutations in that gene might contribute to observed phenotypic changes. Complementing what users see in Gene view, Protein view maps variants onto the protein’s predicted three-dimensional structure, enabling users to visualize clustering within structural motifs and reason about potential impacts on protein folding or function. The color coding is exactly the same in Gene view and Protein view, allowing students to directly see how mutations that occur in distant locations on the gene may even cluster together in three-dimensional space. This provides a concrete framework for discussing how alterations in protein structure can influence folding, stability, and function of the protein. In this way, the combined use of Gene view and Protein is a powerful educational tool—helping students move beyond just identifying gene mutations to reasoning mechanistically about how structural changes drive yeast survivability. In future releases, we plan to expand Protein view to include visualizations of predicted structural changes resulting from mutations to further support hypothesis development. Finally, beyond its use in the classroom, the yEvo Mutation Browser is a tool that can benefit the broader yeast research community by allowing users to upload and compare their data with existing yEvo datasets, enabling quick and easy comparisons to the results of yeast evolution experiments.

A key advantage of the app is its ability to combine data from all classrooms using the same experimental evolution conditions, enabling students to see which of the genes in their groups’ data are recurrently mutated and therefore more likely to be phenotypically causative. Students can also compare mutations observed under one condition to those identified in other evolution conditions, helping them hypothesize about why certain mutations may be beneficial in some environments but not others. For instance, when students explore cumulative data from multiple yEvo experiments within the browser, they might notice that almost all of the conditions lead to mutations in one or more *PDR* genes, particularly *PDR5, PDR3*, and *PDR1*. This observation may prompt students to investigate what makes this gene family so commonly mutated in various environments. Through further exploration, they might learn that the *PDR5* gene encodes a drug transporter, its expression is controlled by the transcription factors Pdr1 and Pdr3, and that variants in all three genes can be involved in adaptive responses to drug exposure (Katzmann *et al*. 1996, Kolaczkowski *et al*. 1996 and 1998). In this example, by identifying the prevalence of mutations in the *PDR* family across multiple conditions, students gain insight into how cells mechanistically respond to different stressors. This serves as a concrete example of how cells utilize drug pumps and related mechanisms to handle diverse environmental challenges, reinforcing broader lessons about cellular adaptation and stress tolerance.

While the yEvo Mutation Browser offers an interactive and user-friendly platform for exploring SNPs and indels in evolved yeast samples, a key limitation is its inability to display other important genomic changes, such as copy number or structural variants (CNVs/SVs). These types of mutations, although significant, have not been included in this initial version of the app. However, there is potential to expand the tool to visualize these mutations in the future, particularly for researchers or advanced students. In published studies using yEvo data, university collaborators have conducted CNV analyses, but these were not a part of classroom discussions. This is a notable limitation, as these mutations often play a critical role in adaptation under specific conditions. For instance, while *PDR5* was frequently mutated in both clotrimazole and caffeine-evolved clones, clotrimazole-evolved samples often involved copy number gains, whereas caffeine-evolved clones had mostly point mutations (Geck *et al*. 2024). These distinctions are currently missing from the Mutation Browser, which focuses solely on SNPs, small indels, and transposon insertions. Nonetheless, the app provides a valuable foundation for visualizing simpler mutation types, and we are actively exploring ways to incorporate CNVs and SVs to provide a more comprehensive view of genomic changes during experimental evolution.

Optimization of the yEvo Mutation Browser is ongoing, based primarily on feedback from users. The reference genome used in the app can also be modified to better represent diverse *S. cerevisiae* strain backgrounds, such as the baking strain used for some prior experiments. For researchers or educators interested in adapting the browser for other strains or species, our detailed GitHub README provides step-by-step instructions. We encourage users to reach out to the authors for assistance if needed.

We believe that yEvo is an incredibly rewarding and effective way to introduce students to scientific inquiry by teaching them microbiology laboratory skills, practicing scientific thinking, and providing an opportunity to participate in real research. The yEvo Mutation Browser enhances this educational experience by providing students with a more meaningful way to visualize and understand the data they generate in the classroom. Through this tool, students can see that while mutations may occur randomly across the genome, those that are frequently selected are most likely driving adaptation. This insight gives students a clearer picture of genetic causality in evolutionary contexts, deepening their understanding of how mutations shape phenotypic outcomes. In conclusion, the yEvo Mutation Browser has allowed us to fill a long-standing gap in our program, bringing the full scope of the yEvo project into clearer focus for students by providing interactive visualizations of their data. As we continue to expand yEvo into additional classrooms, we believe this resource will grow into an invaluable asset for educators, students, and researchers alike, further strengthening the bridge between research and education.

## Data availability

All cumulative yEvo data used in the Mutation Browser as well as the scripts used to generate the app are available at our GitHub repository: https://github.com/dunhamlab/yEvoMutBrowser.git.

## Acknowledgements

We are deeply grateful to Renee Geck for her early input and guidance during the initial brainstorming stages of the project at the 2022 UW Genome Sciences Hackathon, her feedback during development of the app, and her thoughtful review of this manuscript. We also thank all the participants of the 2022 and 2023 GS Hackathon yEvo teams: Nicole Wang (especially for her efforts to rewrite the Python annotation script used in our analysis pipeline), Dennis Godin, Sophie Gibson, Molly Perchlik, Luana Paleologu, Rachel Powell, Conor Camplisson, Matthew Chaw, and Lucas Kampman.

We are also grateful to the yEvo teachers who implemented the app in their classrooms during the 2024-2025 school year and to their students who completed the evolution modules and provided user feedback. In particular, we thank: David Maxwell (The Pingry School), Kelsey Kovarik and Frances Cheong (The Downtown School), Ryan Skophammer (Westridge School), Kenneth Berger, Jonathan Schaper, and Lee Anne Eareckson (Moscow High School), and Michael Delmont (Frontier Senior High School). We are also grateful to Rebecca Brewer and her students at Troy High School for their work on the caffeine evolution experiments, whose data was used for Figure 1.

We acknowledge Sarah McClymont and Brenda Andrews at University of Toronto, along with their undergraduate students, who analyzed over 40 of the evolved samples from Moscow High. Their data was integrated into the cumulative yEvo data presented in the browser. We’d also like to thank Paul Rowley and Claire Warren from University of Idaho for their support in sample curation, phenotype testing, and teacher communication, as well as their continued dedication to the yEvo project as a whole. Finally, we are grateful to M. Bryce Taylor for spearheading the yEvo project at its inception and for generating and analyzing much of the initial data that served as the foundation for the Mutation Browser, and we also appreciate his feedback on the project when the Mutation Browser was first getting started.

---

**Figure S1:**
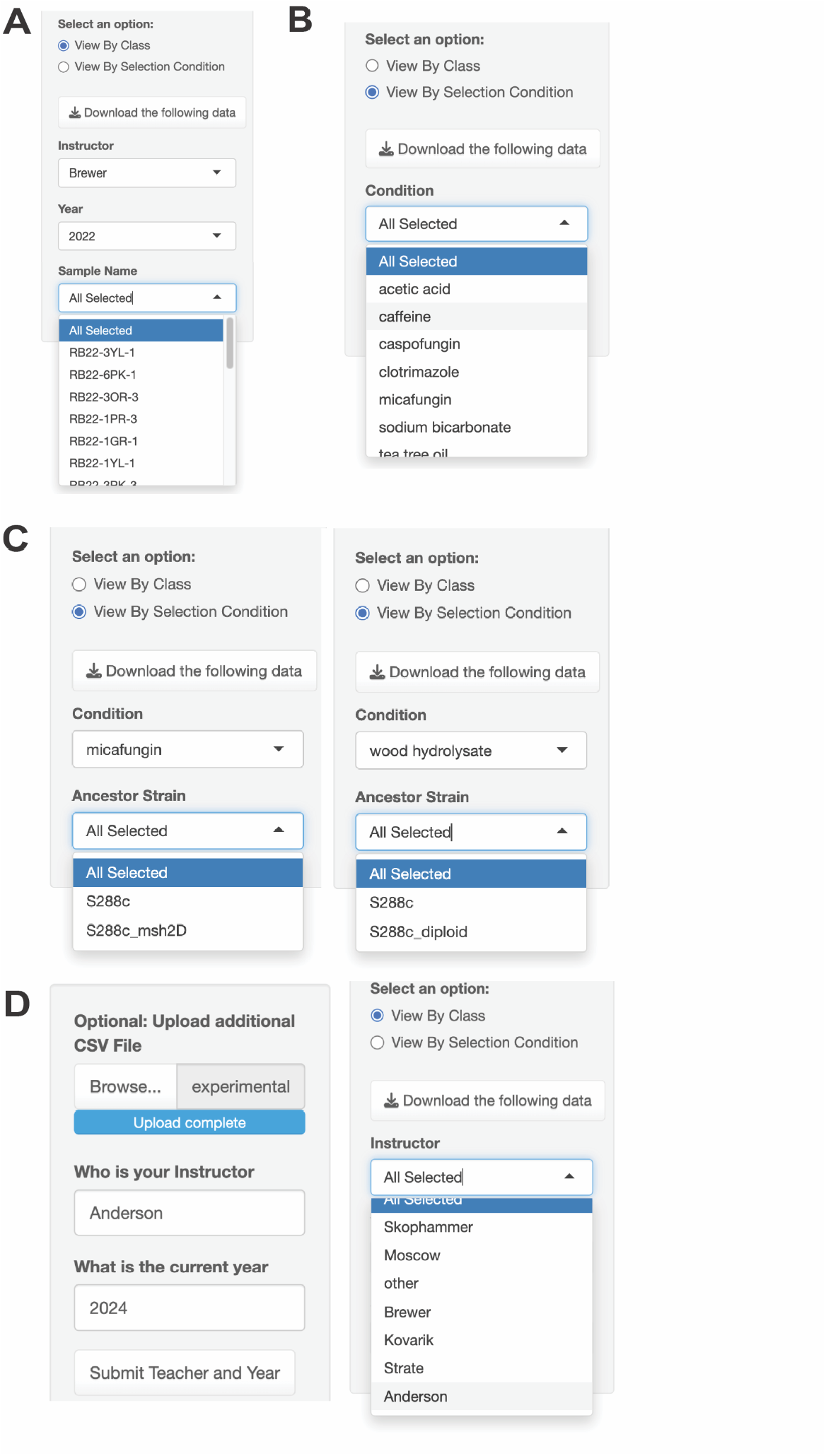
(A) When users select “View By Class”, they will then be provided with dropdown menus for Instructor, Year, and Sample Name. Once Instructor and Year are selected, all samples will be the student groups from that particular classroom. (B) When users select “View By Condition”, all yEvo conditions that have been used for classroom evolution experiments are available in the dropdown menu. Selecting any of these will filter for all yEvo data collected in that condition. (C) Some classrooms used alternative genetic backgrounds of *S. cerevisiae* in their evolution experiments. When this is the case, an additional dropdown menu option will appear, and users can select the desired background. (D) Users also have the option to upload their own CSV of variant data. Once they add their own instructor name and year, the new teacher will appear in the dropdown menu when selecting “View By Class”.

**Figure S2:**
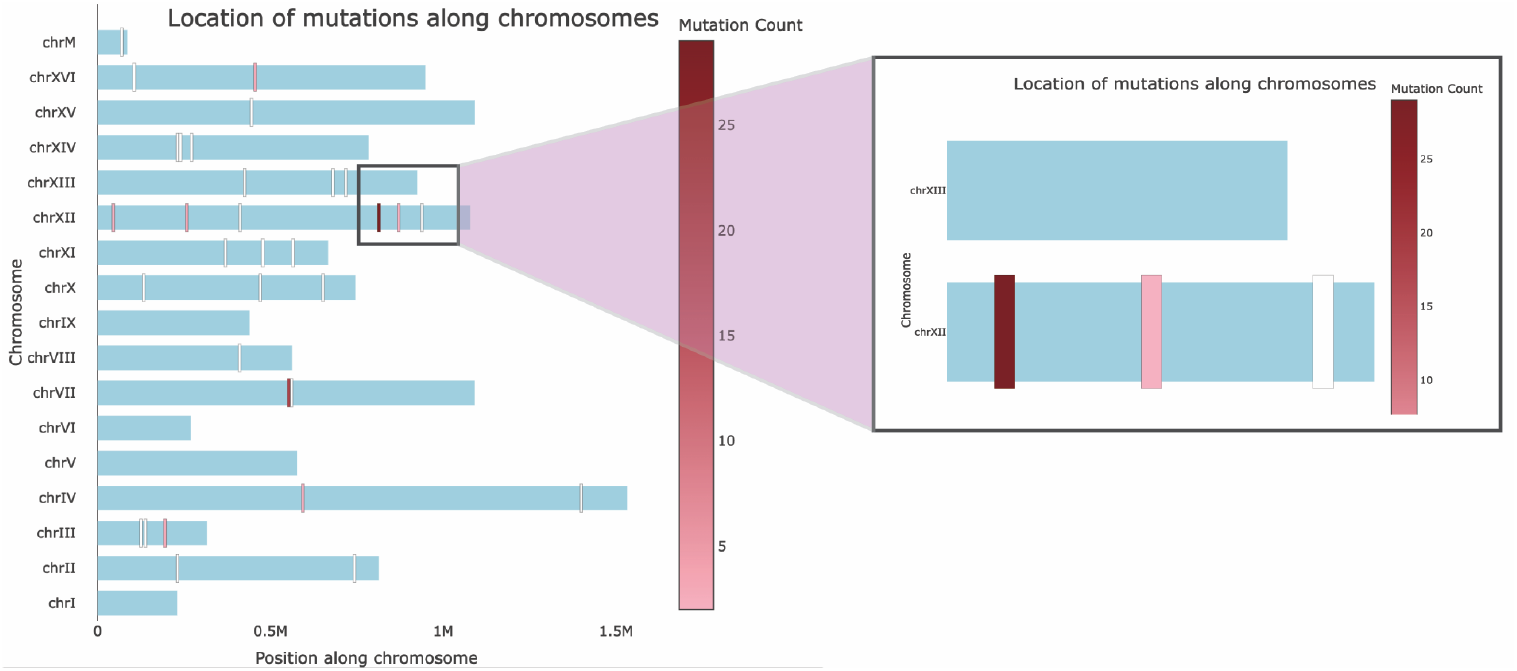
Users can click and drag a box around sites on the chromosome plot to see genes in clustered regions.

**Figure S3:**
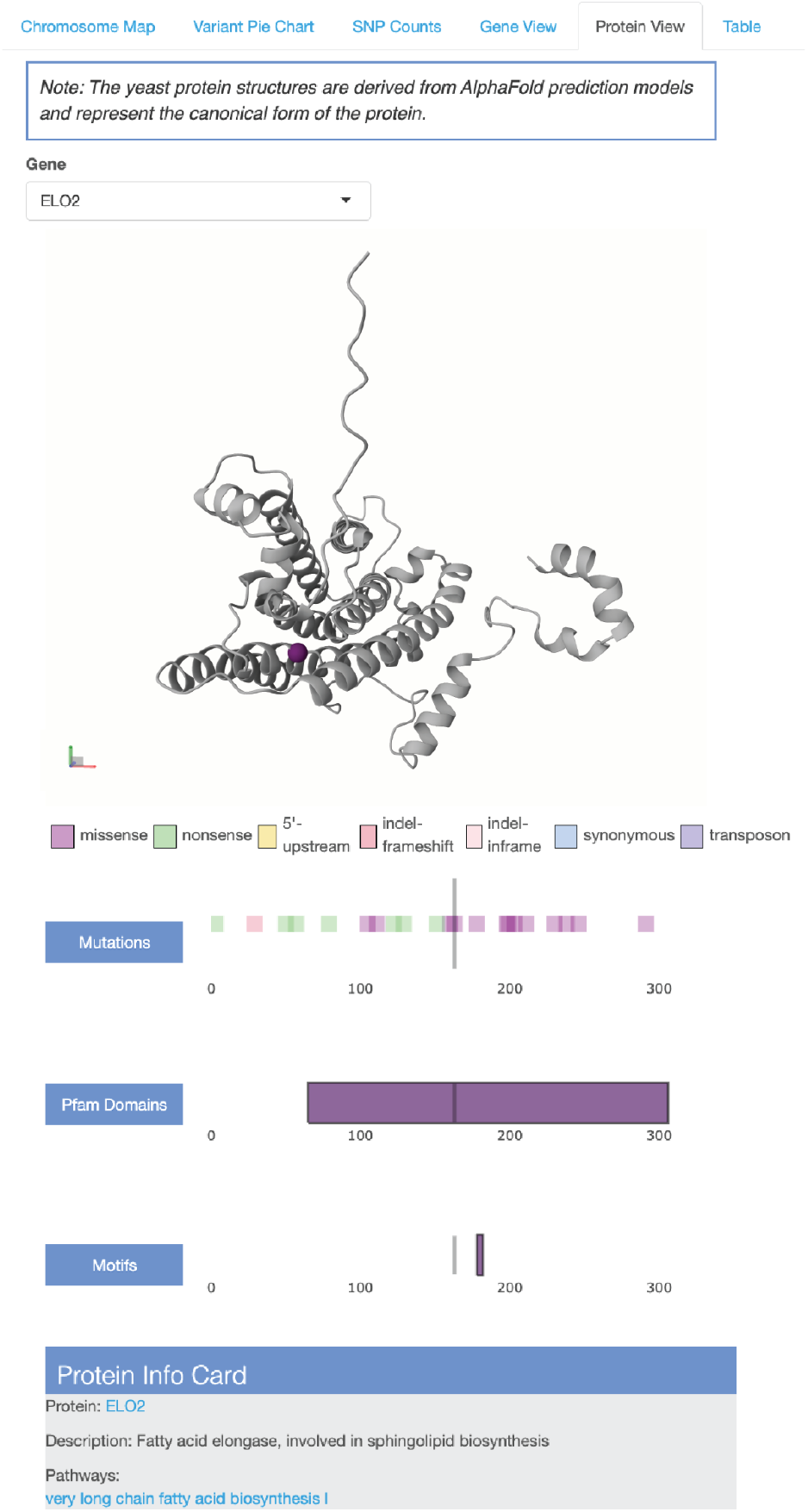
Protein View. Users can select a gene of interest from the dropdown menu to visualize the predicted 3D protein structure encoded by that gene. Linear annotation tracks below the structure display Pfam domains and protein motifs; selecting these features highlights the corresponding regions on the 3D structure. The “Mutations” track indicates the positions of mutations present in the selected yEvo dataset along the linear amino acid sequence, color coded by mutation type in accordance with the Gene View plots. Mutations may be selected individually or in aggregate to visualize their localization on the protein structure.

